# Delta oscillations coordinate intra-cerebellar and cerebello-hippocampal network dynamics during sleep

**DOI:** 10.1101/2021.05.04.442571

**Authors:** A Torres-Herraez, TC Watson, L Rondi-Reig

## Abstract

During sleep, the widespread coordination of neuronal oscillations across both cortical and subcortical brain regions is thought to support various physiological functions, including memory consolidation. However, how sleep-related activity within the brain’s largest sensorimotor structure, the cerebellum, is multiplexed with well described sleep-related mechanisms in regions such as the hippocampus remains unknown.

To address this gap in knowledge, we simultaneously recorded from the dorsal hippocampus and three distinct regions of the cerebellum (Crus I, lobule VI and lobules II/III) during natural murine sleep. We found that LFP oscillations are coordinated between the two structures in a sleep-stage specific manner. Particularly during non-REM sleep, prominent delta frequency coherence was observed between lobule VI and hippocampus. We additionally observed that non-REM associated hippocampal sharp wave ripple activity can drive discrete LFP modulation in all the recorded cerebellar regions, with the shortest latency effects observed in lobule VI.

We also describe discrete phasic sharp potentials, synchronized across cerebellar regions, which were strongly phase locked to the peak of ongoing cerebellar delta oscillations and found in greatest numbers during REM. These phasic sharp potentials recorded within the cerebellar cortex were found to be phase-locked to the trough of the hippocampal theta oscillation, further suggesting cross-structural coordination. During REM, cerebellar delta oscillation phase significantly modulated hippocampal theta frequency, and this effect was greatest when phasic sharp potentials were most abundant. Within all three cerebellar regions, prominent LFP oscillations were observed at both low (delta, <4 Hz) and very high frequencies (~ 250 Hz) during non-REM and REM sleep. Intra-cerebellar cross-frequency analysis revealed that delta frequency oscillations strongly modulate those in the very high frequency range.

Together, these results reveal multiple candidate physiological mechanisms to support ‘offline’, bi-directional interaction within distributed cerebello-hippocampal networks. In particular, we describe a prominent cerebellar delta oscillation, which appears to act as a temporal coordinator of cerebellar activity at both local (intra-cerebellar) and distributed (cerebello-hippocampal) network levels.

## Introduction

Many important physiological processes are associated with sleep, including memory formation and consolidation (Diekelmann and Born, 2010). In particular, sleep-related neurophysiological events in the hippocampus, such as place cell reactivations and high frequency sharp wave ripples are vital to spatial memory formation (e.g. de Lavilléon et al., 2015). Additionally, widespread slow wave oscillations are thought to play a crucial role in the temporal coordination of discrete sleep-related events, such as hippocampal ripples and thalamo-cortical spindles, across a large network of brain regions (e.g. Nicola et al., 2019)

Recent studies have shown that during wakeful behaviour the cerebellum is crucial for hippocampal place cell stability and efficient navigation (Rochefort et al., 2011; Lefort et al., 2019; Burguière et al., 2005, Babayan et al., 2017). Furthermore, the neuroanatomical connectivity required to support this influence has been recently described and physiological correlates of cerebello-hippocampal interaction identified ‘online’ during goal-directed navigation (Watson et al., 2019). So far, however, nothing is known about possible ‘offline’ cerebello-hippocampal physiological interactions during sleep.

To date, sleep research has primarily focused on state-dependent correlation in neocortical and subcortical structures. However, early studies in epileptic patients implanted with intra-cerebellar electrodes (Niedermeyer and Uematsu, 1974) and recent work using non-invasive techniques (Jahnke et al., 2012; Kaufmann et al., 2006; Schabus et al., 2007) have described state-dependent modulation of cerebellar activity during sleep in humans. During non-rapid eye movement (non-REM) sleep, coordinated slow oscillations, recorded predominantly in cerebellar vermis and fastigial nucleus (Niedermeyer and Uematsu, 1974; Schabus et al., 2007), are correlated with neocortical K-complexes (Jahnke et al., 2012) and sleep spindles (Schabus et al., 2007). During REM sleep, an increase in blood-oxygen-level-dependent signals has been observed in both the cerebellar vermis and hemispheres (Braun, 1997).

Similarly, single-cell recording in animal models has revealed sleep-state related changes in both Purkinje cell and deep cerebellar nuclei neuron firing. During non-REM, no clear neuronal firing rate changes were observed either in cats (Hobson and McCarley, 1972; Marchesi and Strata, 1970; McCarley and Hobson, 1972; Palmer, 1979) or monkeys (Mano, 1970). However, a reduced probability for short interspike intervals was observed in Purkinje cell simple spike firing (McCarley and Hobson, 1972). Consistent with observations made in humans, during REM sleep a significant increase in the firing rate of Purkinje cells was observed in the cerebellar cortex of cats (Hobson and McCarley, 1972; Marchesi and Strata, 1970) and monkeys (Mano, 1970). The presence of large amplitude, phasic events in the cerebellum of cats (Harlay et al., 1974; Pellet and Harley, 1977) and rats (Marks et al., 1980), mainly during REM sleep epochs, has also been attributed to the transmission of ponto-geniculo-occipital waves (PGO waves; Farber et al., 1980; Velluti et al., 1985). Finally, recent studies have highlighted cerebellar roles in the generation of sleep spindles in monkeys (Xu et al., 2020) and sleep-wake transitions in mice (Zhang et al., 2020).

Thus, in addition to the multitude of studies highlighting the functional importance of sleep-state dependent processes in the hippocampus (see Klinzing et al., 2019 for review) the aforementioned evidence indicates clear cerebellar activity modulation during sleep (reviewed in Canto et al., 2017). Nevertheless, our understanding of spatial and temporal organisation of cerebellar activity across sleep states remains rudimentary. Furthermore, to the best of our knowledge, nothing is known about the physiological links between cerebellum and hippocampus during sleep. We therefore simultaneously recorded spontaneous local field potentials (LFPs) from the dorsal hippocampus, vermal cerebellar lobules II/III (Lob II/III), VI (Lob VI) and hemispheric Crus I in freely behaving and sleeping mice. Our analysis reveals that delta oscillations, along with associated phasic sharp potentials (PSPs), coordinate both local cerebellar and distributed cerebello-hippocampal network dynamics in a sleep-state dependent manner.

## Results

### Spectral profiling of cerebellar and hippocampal activity across sleep states

In line with previous studies (e.g. Montgomery et al., 2008), hippocampal LFP was dominated by sustained, low amplitude theta oscillations (6 - 12 Hz) during wake and REM states, and high amplitude slow oscillations in the delta range (< 4 Hz) during non-REM sleep (Figure 1 A-C).

**Figure 1.**
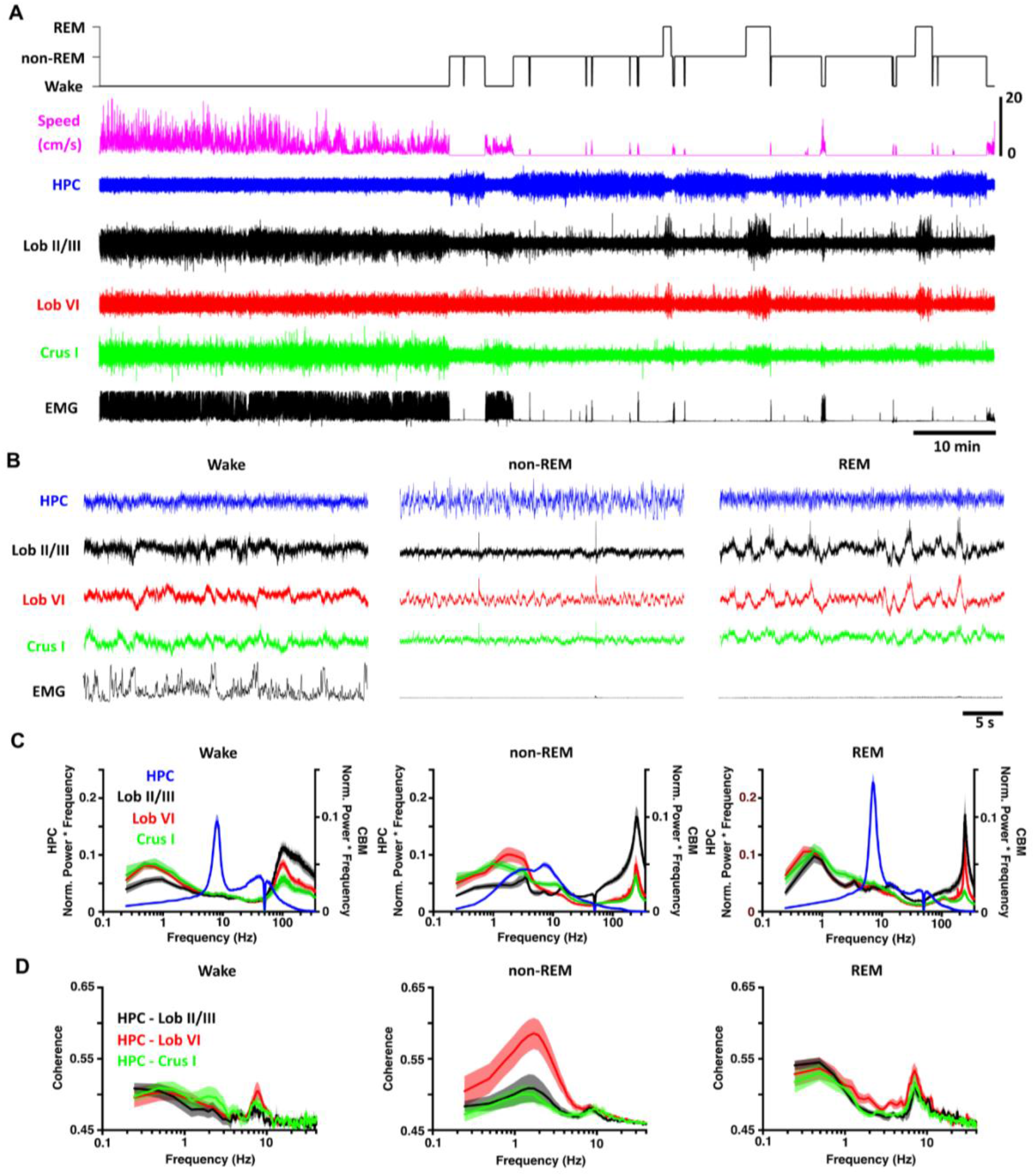
Sleep-state specific activity patterns are present in the cerebello-hippocampal network. **A,** Hippocampal (HPC) and cerebellar cortical (Crus I, Lob VI and II/III) LFPs were recorded as mice cycled between defined wake, non-REM and REM states. **B,** During wake, hippocampal theta and cerebellar < 1 Hz and high gamma (100 - 160 Hz) oscillations occurred concomitantly. Similarly, during REM, hippocampal theta oscillations were accompanied by widespread delta (< 4 Hz) and very fast (~ 250 Hz) cerebellar oscillations. During non-REM, high amplitude hippocampal activity co-occurred with both slow, phasic and very high frequency (~ 250 Hz) cerebellar oscillations. **C,** Mean power spectra for each of the defined states (HPC, n = 20; Crus I, n = 11; Lob VI, n = 15; Lob II/III, n = 11). **D,** Mean coherence between HPC-Crus I (n = 11); HPC-Lob VI (n = 15) and HPC-Lob II/III (n = 11) across states.

Conversely, high amplitude low frequency oscillations (including < 4 Hz delta waves) were characteristic of wake, REM and non-REM states in the cerebellum (Figure 1 A-C). However, differences in delta oscillation power, as well as peak frequency, were observed both across cerebellar subregions and wake/sleep states (Figure 1C, Supplemental Figure 1A-B). Similarly, a significant shift in the peak delta frequency was observed during non-REM sleep, specifically in Lob VI (Supplemental Figure 1B). In contrast, these differences were not observed during REM, when similar spectral profiles were found across cerebellar regions (Figure 1C). Moreover, delta oscillations were highly coherent between cerebellar recording sites (Figure 1B, Supplementary Figure 2), suggesting they may play a role in intra-cerebellar coordination during REM sleep.

In addition to delta activity during wake, high-frequency oscillations (100-160 Hz) were also observed in the cerebellar power spectra (Figure 1C). However, during both non-REM and REM states, power in the high-frequency range was overshadowed by a prominent, narrowband ~240-280 Hz oscillation (classified as very-high frequency oscillations, VHFOs; de Solages et al., 2008; Figure 1C; Supplemental Figure 1C). Local variability within this frequency band was observed across cerebellar regions and sleep states, with maximal values observed in Lob II/III during non-REM and REM sleep (Supplemental Figure 1D). In contrast, Crus I VHFO power was particularly reduced in REM (Figure 1C, Supplemental Figure 1D, paired t-tests: Crus I: n = 11, p = 0.0005; Lob II/III, n = 11, Crus I, n = 11; Lob VI, n = 15; Lob II/III, n = 11 mice).

Overall, these data illustrate the presence of lobule and sleep-state dependent LFP dynamics across anterior (Lob II/III), posterior (Lob VI) and hemispheric (Crus I) cerebellum and highlight the existence of a prominent cerebellar delta oscillation.

### Cerebello-hippocampal coherence across sleep states

Next, we sought to measure the level of functional coupling (measured as coherence; cf Watson et al., 2019; Xu et al., 2020) between the cerebellum and hippocampus during wake/sleep states (Figure 1D, E). During wake, coherence peaks were similarly observed across delta and theta frequencies in all cerebello-hippocampal combinations (Figure 1D, left panel). Strikingly, as mice moved to non-REM sleep, delta band coherence between hippocampus and Lob VI was significantly increased relative to other recorded cerebellar regions (Figure 1D, Supplemental Figure 3) indicating region-specific cerebello-hippocampal coupling during this sleep state. Furthermore, significant modulation of delta and theta coherence across sleep states was present in HPC-Lob VI and HPC-Lob II/III recording combinations, while remaining stable between HPC-Crus I (Figure 1D; Supplemental Figure 3). In contrast, during REM sleep, coherence patterns were similar to wake, with the most prominent peaks observed in the theta and low frequency range (< 1Hz; Figure 1D, right panel; Supplemental Figure 3).

In summary, by characterizing the spectral properties of cerebello-hippocampal network LFP during wake and sleep we reveal that coherence between these regions is sleep-state dependent and particularly prominent between hippocampus and Lob VI during the non-REM stage of sleep.

### REM associated cerebellar phasic sharp potentials phase lock to both local cerebellar and distant hippocampal oscillations, respectively

In addition to the striking sleep-state related changes observed in ongoing cerebellar LFP oscillations, we also noted prominent phasic sharp potential events (PSPs; large-amplitude voltage fluctuations of 129.5 ± 5.68 ms duration and 2.31 ± 0.15 z-score amplitude) across the three cerebellar recording sites (Figure 2A-C). PSPs occurred in all three cerebellar regions at similar times (cross-correlation between all pairs of cerebellar recorded PSPs revealed a peak at 0 latency; Figure 2A-C) and were found during both REM and non-REM epochs; however, they occurred in greatest numbers during REM (PSP density during non-REM = 0.12 ± 0.01, PSP density during REM = 0.5 ± 0.05, Wilcoxon test P < 0.0001; Figure 2A, D). Moreover, PSPs occurred in concentrated clusters during REM (inter-event interval < 1s) but were distributed more sparsely during non-REM (Figure 2E). We also observed that during REM epochs, PSPs tended to occur at specific phases of the ongoing cerebellar delta waves in all three cerebellar regions (Figure 2A). This observation was confirmed by computing the distribution of PSP events relative to the phase of REM associated cerebellar delta waves (Figure 2F-H; PSPs occurred most often near the peak of the cycle with a preferred angle of 350.6 ± 2.248 ° and with similar levels across all three cerebellar regions, Figure 2G, H). Next, we investigated phase locking of PSPs recorded in the cerebellum relative to hippocampal LFP theta phase during REM (Figure 2I), which revealed that these events occur consistently near the trough of the theta cycle (preferred phase angle, 147.4 ± 15.93 °; Figure 2I) and at similar levels across cerebellar regions (Figure 2 J,K).

**Figure 2.**
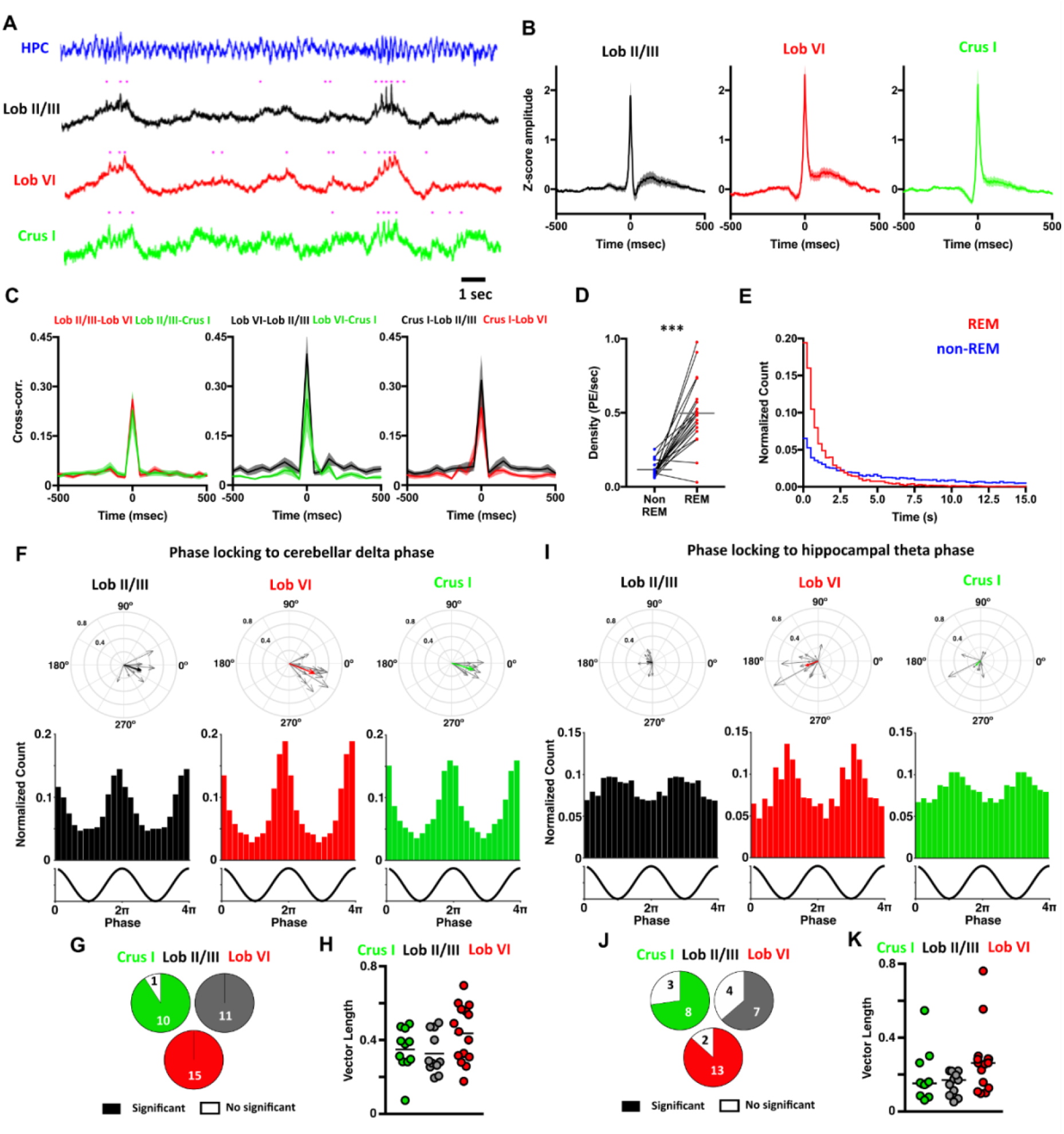
Phasic potentials (PSPs) occur at preferred phases of cerebellar delta and hippocampal theta oscillations during REM. **A,** We observed phasic potentials in the cerebellar LFP recordings during REM (indicated by purple dots), which occurred preferentially during up-phase of the delta oscillations. Moreover, hippocampal theta frequency was transiently accelerated during the up-phase of the cerebellar delta oscillations. **B,** PSPs were found in all cerebellar recording sites and all mice. Averaged waveforms across animals (Lob II/III n = 11; Lob VI n = 15; Crus I n = 11). **C,** Cross-correlograms between the PSPs detected at a given cerebellar region and those recorded in the others. All cross-correlograms show a peak with ≤ 50ms lag suggesting that most of the PSPs co-occurred at the three cerebellar regions near simultaneously. **D,** PSPs were observed during non-REM and REM sleep, however, the density of these events was significantly higher during REM (non-REM density = 0.1152 ± 0.012 PSPs/s; REM density = 0.4973 ± 0.050 PSPs/s; paired t-test, P < 0.0001). **E,** The distributions of the inter-event intervals (IEIs) were significantly different between non-REM and REM epochs (Two-sample Kolmogorov-Smirnov test, P < 0.0001). During REM, PSPs were preferentially concentrated as clusters (IEIs < 1 s). **F,** PSPs recorded at all cerebellar regions were phase-locked to the local delta oscillations during REM. In the top panels, the individual vectors of the phase locking for each mouse (grey arrows) and the average vector across all mice (colour coded; resultant vector angle: Crus I = 342.97 °, Lob II = 343.05 °, Lob VI = 338.67 °. Resultant vector length: Crus I = 0.34, Lob II = 0.28, Lob VI = 0.41). In the bottom panels, the normalized count of PSPs recorded at the different phases of delta oscillations in all mice for each cerebellar region (Lob II/III = 4004 PSPs/11 mice, Rayleigh’s test, P < 0.001; Lob VI = 3445 PSPs/15 mice, Rayleigh’s test, P < 0.001; Crus I = 3398 PSPs/11 mice, Rayleigh’s test, P < 0.001). **G**, Fraction of mice with significant phase locking (significant phase locking was found in Crus I, 10/11 mice; Lob II/III, 11/11 mice; Lob VI 15/15 mice). **H**, The level of the phase locking did not differ across cerebellar regions (1-way ANOVA, P = 0.0932). **I-K**, Same as for **F-H** but here calculating phase locking of PSPs detected in the cerebellum to ongoing hippocampal theta oscillations. **I**, Top row; Lob II/III Rayleigh’s test, P < 0.001; Lob VI Rayleigh’s test, P < 0.001; Crus I Rayleigh’s test, P < 0.001. **J**, Fraction of mice with significant phase locking of PSPS to hippocampal theta (significant phase locking was found in Crus I, 8/11 mice; Lob II/III, 7/11 mice; Lob VI 13/15 mice). **K**, The level of the phase locking to hippocampal theta did not differ across cerebellar regions (Kruskal-Wallis Test of vector lengths, P = 0.0717).

### Intracerebellar cross-frequency modulation: cerebellar delta is a modulator of VHFOs during REM

The most prominent, ongoing LFP oscillations observed in the cerebellum during REM sleep were found in the delta (<4 HZ) and VHFO (~ 250Hz) frequency ranges, respectively (see Figure 1). Thus, to investigate any relationship between these two oscillators, we conducted cross-frequency spectral analysis (Figure 3). This revealed that during REM, the power of cerebellar VHFOs is significantly modulated by the phase of those in the delta range (Figure 3B, C). Interestingly, this modulation was significant only in vermal regions (Lob VI, II/III) and not in Crus I. This finding is consistent with the reduced presence of VHFOs during REM in this cerebellar region compared to the others recorded (Figure 1C, Supplemental Figure 1D), further indicating spatial heterogeneity in cerebellar processing during sleep.

**Figure 3.**
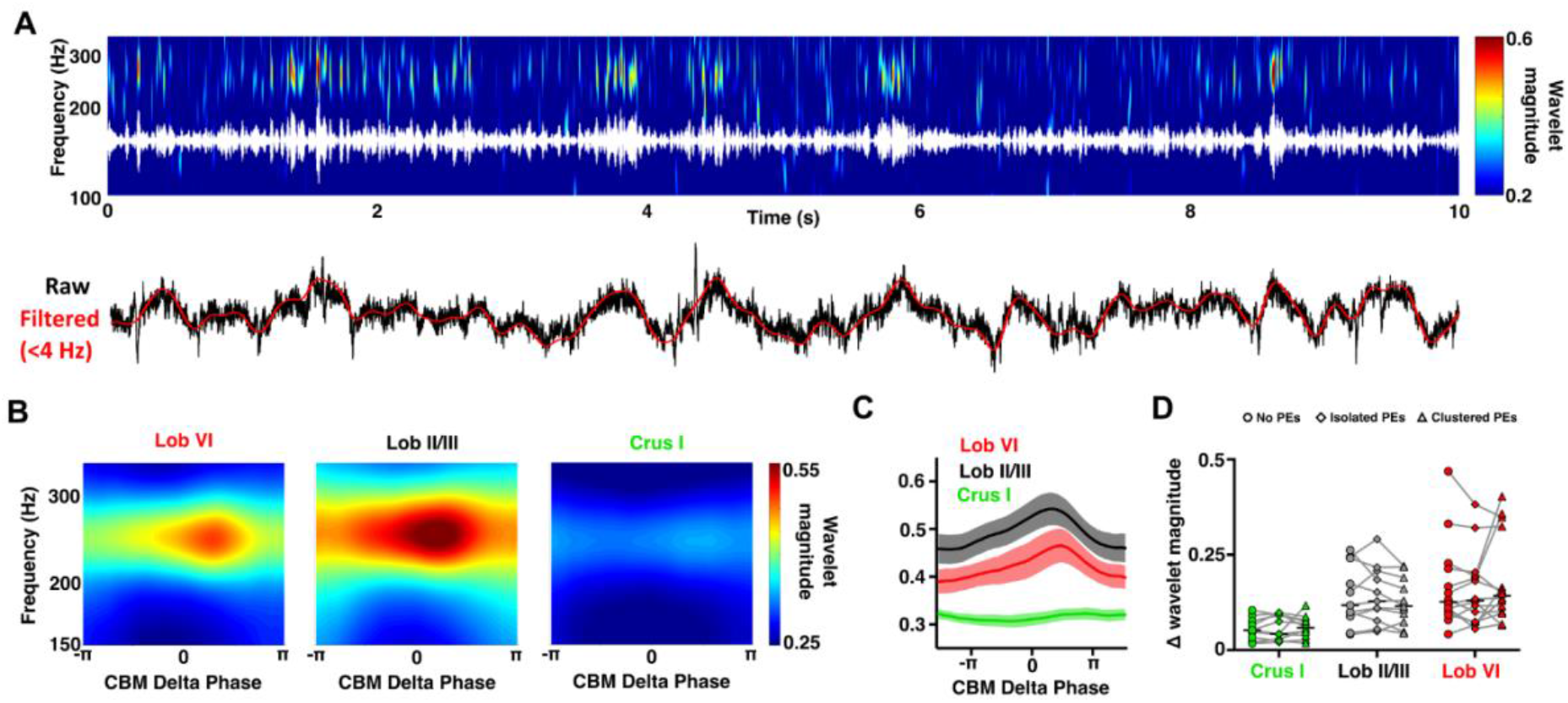
Modulation of VHFOs frequency oscillations by delta oscillations within the cerebellum during REM. **A**, Representative spectrogram (top; with VHFO frequency filtered trace overlaid in white) and raw cerebellar LFP recordings (bottom; with delta frequency filtered oscillations overlaid in red) showing the modulation of VHFOs by cerebellar delta oscillations during a REM epoch. The cerebellar LFP is dominated by both delta oscillations and VHFOs, which fluctuate dynamically within the ~250Hz and <4 Hz ranges, respectively. The fluctuations in cerebellar VHFO power appear temporally aligned to the prominent delta oscillations in the cerebellar LFP. **B**, Average spectrograms aligned to the phase of the cerebellar delta oscillations (<4 Hz) during REM sleep. The power of cerebellar VHFOs appears greatest following the peak of cerebellar delta (zero phase radians) in vermal lobules (VI, II/III) but not Crus I. **C**, Average cerebellar VHFO power aligned to cerebellar delta oscillation phase (RM 2-Way ANOVA Cerebellar Region x Delta Phase: Delta Phase P < 0.0001, Cerebellar region P = 0.0001, Interaction P = 0.0002. Multiple comparisons of power across delta phase compared to trough value with FDR correction: Crus I: p>0.05, Lob II: p < 0.01 from −18.9 ° - 78.5 ° (−0.105 π - 0.44 π) Lob VI: p < 0.01 from −3.72 ° - 93.97 ° (−0.0 2π-0.52 π)). **D**, Effect of PSP abundance during the cerebellar delta oscillations on the modulation of VHFOs during REM. We compared the maximal change in cerebellar VHFO power during cerebellar delta oscillations in which no PSPs, isolated PSPs or clusters of PSPs were detected. No significant differences were observed for the three cerebellar regions (2-Way ANOVA Cerebellar Region x Content of PSPs: Content of PSPs P = 0.8449, Cerebellar Region P = 0.003, Interaction P = 0.2601, Multiple comparisons across cerebellar region effect with FDR correction: Crus I x Lob II, corrected p = 0.0101, Crus I x Lob VI, corrected p = 0.0009, Lob II x Lob VI, corrected p = 0.1004). In all mean plots, n = Lob VI, n = 15 mice; Crus I, n = 11 mice; Lob II/III, n = 11 mice.

Since PSPs were highly abundant during REM and also phase locked to cerebellar delta, we investigated whether they play a role in local cross-frequency modulation between delta phase and VHFO power. By comparing modulation in delta cycles with no PSPs, those containing isolated PSPs and those containing clusters of PSPs (Figure 3D), we found that PSP density did not significantly influence the degree of cross-frequency modulation observed in any of the recorded cerebellar regions (Figure 3D).

### Cerebellar delta and phasic sharp potentials modulate hippocampal theta oscillations during REM

We next investigated if the prominent cerebellar delta oscillations and the high density of PSPs observed during REM was related to, or impacted upon, theta oscillations in the hippocampus, which are a well described physiological signature of REM sleep (Figure 4). During REM, hippocampal theta oscillations fluctuated within a frequency range of approximately 6-12 Hz and visual inspection of spectra and raw LFP traces revealed that these theta frequency fluctuations appeared to be temporally aligned with cerebellar delta oscillation cycles/PSP events (Figure 4A). Therefore, we next calculated hippocampal LFP power spectra triggered from the peak of cerebellar delta wave cycles. This analysis revealed that hippocampal theta frequency increased during the ascending phase of the cerebellar delta oscillation when compared with the values at the trough (-π) (triggered from Crus I: maximal frequency of hippocampal theta = 7.97 ± 0.095 Hz at −18.74 ° (−0.1 π); triggered from Lob II: maximal frequency of hippocampal theta = 7.94 ± 0.095 Hz at −3.72 ° (−0.02 π); triggered from Lob VI: maximal frequency of hippocampal theta = 8.14 ± 0.154 Hz at −18.74 ° (−0.1 π) Figure 4B,C).

**Figure 4.**
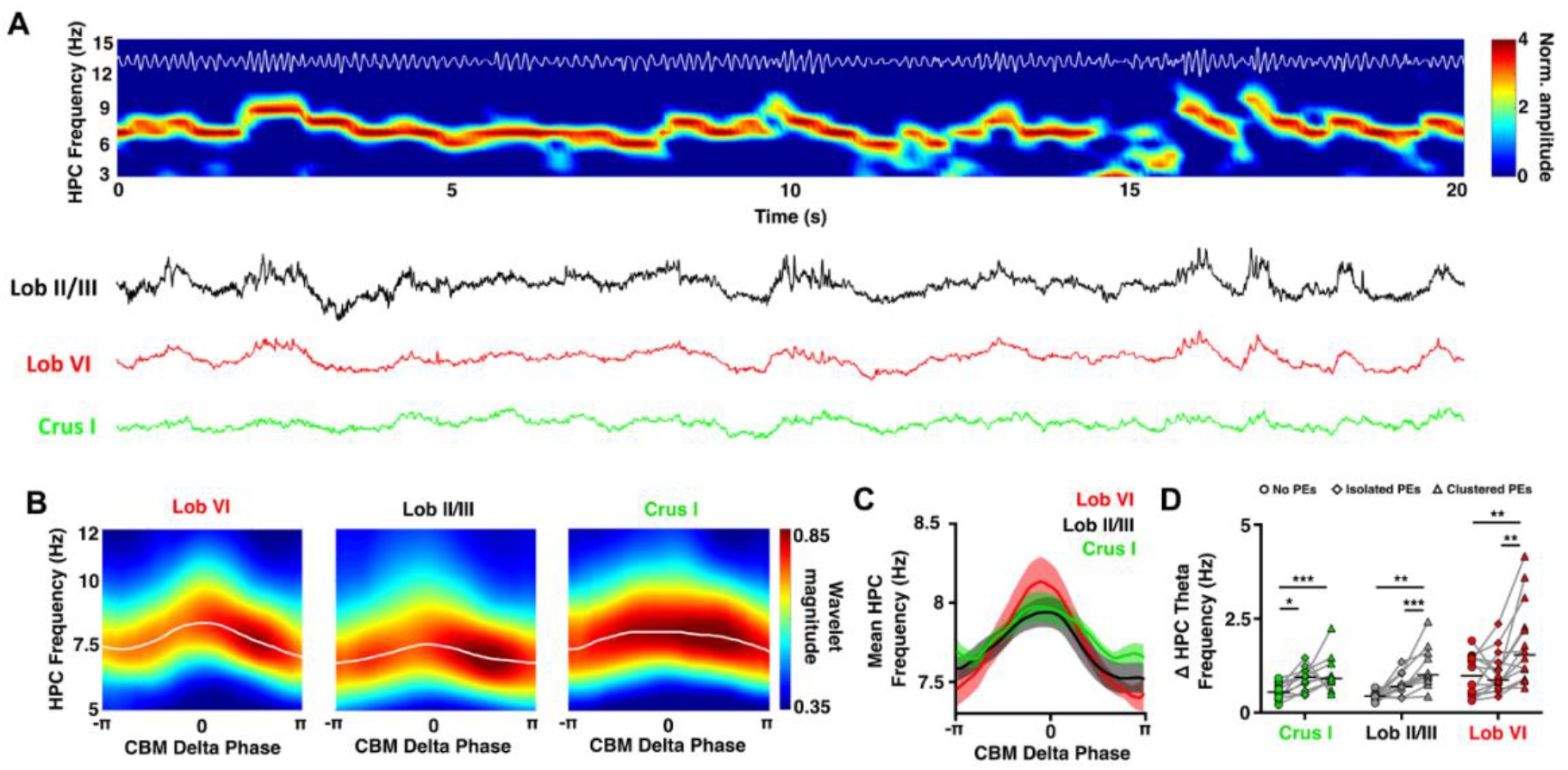
Modulation of hippocampal theta frequency by cerebellar delta oscillations during REM. **A**, Representative spectrogram (top; with theta filtered trace overlaid in white) and raw cerebellar LFP recordings (bottom) showing the modulation of hippocampal theta frequency by cerebellar delta oscillations during a REM epoch. The hippocampal LFP is dominated by theta oscillations, which fluctuate dynamically within the 6-12 Hz range. These fluctuations in hippocampal theta frequency appear temporally aligned to the delta oscillations dominating cerebellar LFPs (examples of this are indicated by black horizontal bars above spectrogram). **B**, Example hippocampal spectrograms aligned to the phase of the cerebellar delta oscillations (<4 Hz) during REM sleep. The preferred hippocampal theta frequency (white line, overlaid) shows an acceleration coincident with the peak of the cerebellar delta oscillations (0 phase radians). **C**, Average preferred hippocampal theta frequency aligned to cerebellar delta oscillation phase (Lob VI, n = 15; Crus I, n = 11; Lob II/III, n = 11). Significant hippocampal theta frequency acceleration was found from the trough of delta to the peak of delta in all cerebellar regions. However, no differences in hippocampal theta frequency modulation were observed between the delta oscillations recorded across the three cerebellar locations (repeated measurements Two-way ANOVA phase x cerebellar region, phase effect p < 0.0001, region effect p = 0.7953, interaction effect p = 0.0295). **D**, Effect of PSP abundance during the cerebellar delta oscillations on the modulation of preferred hippocampal theta frequency during REM. We compared the maximal change in hippocampal theta frequency during cerebellar delta oscillations in which no PSPs, isolated PSPs or clusters of PSPs were detected. Significant differences were observed for the three cerebellar regions (Crus I, Friedman test with FDR correction, p = 0.0011; Lob II/III, Friedman test with FDR correction, p < 0.0001; Lob VI, Friedman test with FDR correction, p = 0.0015) with a significant increase in the hippocampal theta modulation during delta waves when clusters of PSPs were detected compared with those with no PSPs or isolated PSPs.

Given that PSP timing is locked to specific phases of both cerebellar delta and hippocampal theta oscillation cycles (Figure 2), we next investigated if the number of PSPs occurring during the cerebellar delta wave was associated with the level of hippocampal theta modulation. In contrast to intra-cerebellar delta phase modulation of VHFOs, by comparing cycles with no PSPs, those containing isolated PSPs and those containing clusters of PSPs (Figure 4D), we found that the acceleration of hippocampal theta was most prominent when clusters of cerebellar PSPs were present (PSP quantity x cerebellar < 4 Hz phase repeated measures two-way ANOVA, quantity of PSP effect, F(1,4) = 43.94, p = 0.0027; interaction effect F(47,188) = 4.678, p-value < 0.0001), suggesting that these phasic events are linked to the observed changes in hippocampal theta frequency.

### Hippocampal ripples drive LFP changes in the cerebellum

During REM, dominant hippocampal theta oscillations are coordinated with prominent cerebellar LFP network activity (delta oscillations and PSPs). During non-REM sleep, however, hippocampal activity is characterized by the presence of prominent sharp-wave ripples (SWRs). Therefore, we next asked if cerebellar activity is coordinated or modulated by hippocampal SWRs during non-REM sleep. To address this question, we first detected SWRs (Figure 5A) and computed the cerebellar LFP averaged relative to SWR maximal amplitude (Figure 5B). Event related field potentials (ERPs) were clearly observed in all the recorded cerebellar regions. The SWR triggered responses were characterized by a prominent slow component that preceded the onset timing of SWR maximal amplitude followed by an ERP (occurring after SWR ripple maximal amplitude; Figure 5B). In order to compare the amplitude of the ERPs across cerebellar regions, we normalized them by subtracting the pre-SWR maximal baseline levels (Figure 5B inset). The onset-to-peak amplitude in normalized ERPs did not differ across cerebellar regions (Figure 5C, left panel). In contrast, latency to ERP peak was shortest in Lob VI (32 +- 7.89 ms) compared to Crus I and to Lob II/III (Figure 5C, right panel).

**Figure 5.**
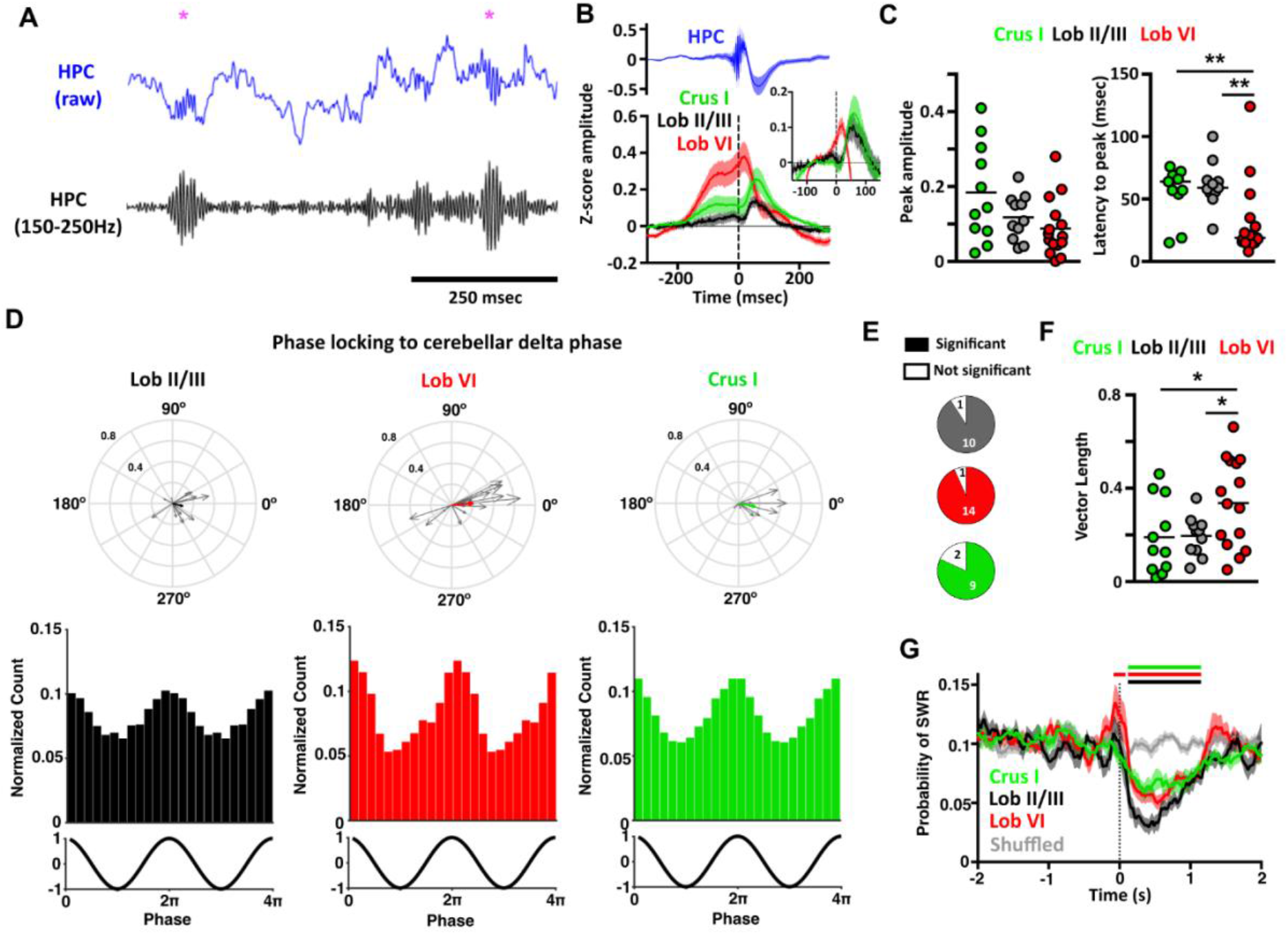
Hippocampal sharp-waves ripples (SWR) trigger evoked activity in the cerebellum. **A,** Example of the raw (blue) and filtered (150-250 Hz, black) hippocampal LFP during non-REM sleep. SWR were identified (purple asterisks) at the maximal value in the filtered signal. **B,** Averaged SWR-triggered waveforms from the cerebellar recordings. Zero time represents the time of detected SWR peak in the filtered signal. Evoked field potentials were observed in all the cerebellar recordings. The averaged waveforms in Lob VI and Crus I revealed that the SWR occurred at the peak of cerebellar delta oscillations. Inset, shows the same data but baselined against values from −100 to −50 msec, eliminating the slow frequency component and allowing clearer visualization of the evoked potentials. **C,** From the baseline normalised data, we quantified the ERP amplitude measured as the onset-to-peak value and also the onset latency. No significant differences were observed for amplitude (Crus I, mean amplitude = 0.184 +- 0.04; Lob II, mean amplitude = 0.118 +- 0.02; Lob VI, mean amplitude = 0.088 +- 0.02; Kruskal-Wallis test P = 0.0968); however, Lob VI ERPs occurred at significantly shorter latencies compared to both Lob II/III and (Crus I: mean = 55.91 +- 6.14 ms; Lob II/II: mean = 61.45 +- 5.53 ms; Lob VI: mean = 32 +- 7.89 ms; Kruskal-Wallis test P = 0.0053; Multiple comparisons FDR corrected: Lob VI vs Crus I P= 0.0044, Lob VI vs Lob II P= 0.0044, Crus I vs Lob II P = 0.3034). **D,** Significant phase locking of SWRs to delta oscillations was found in all cerebellar regions during non-REM sleep. In the upper panels, the normalized count of PSPs recorded at the different phases of delta oscillations in all mice for each cerebellar region (Lob II/III = 14012 SWRs/11 mice, Rayleigh’s test, P < 0.001; Lob VI = 11915 SWRs/15 mice, Rayleigh’s test, P < 0.001; Crus I = 15532 SWRs/11 mice, Rayleigh’s test, P < 0.001). In the lower panels, the individual vectors of the phase locking for each mouse (grey arrows) and the average vector across all mice (colour coded; resultant vector angle: Crus I = 348.27 °, Lob II = 345.4 °, Lob VI = 4.26 °. Resultant vector length: Crus I = 0.16, Lob II = 0.1, Lob VI = 0.21). **E**, Fraction of mice with significant SWR-to-cerebellar delta phase locking (significant phase locking was found in Crus I, 9/11 mice; Lob II/III, 10/11 mice; Lob VI 14/15 mice). **F,** The level of SWR phase locking was higher with Lob VI compared to Crus I or Lob II/III (1-way ANOVA, P = 0.0324; multiple comparisons FDR corrected, Lob VI-Crus I P = 0.0149, Lob VI-Lob II/III P = 0.0149, Crus I-Lob II/III P = 0.3317). **G**, Probability of SWRS relative to PSP onset. The number of SWRs are reduced following PSP occurrence (at time zero; cf. Tsunematsu et al., 2020; 2-way ANOVA, region vs time, time effect P < 0.0001, region effect P < 0.0001, interaction effect P < 0.0001. Multiple comparisons between cerebellar regions and shuffled data). For all mean plots, Crus 1, n = 11 mice; Lob II/III, n = 11 mice; Lob VI, n = 15 mice.

We next asked whether the slow frequency component observed in the SWR-triggered responses resulted from the timing of SWRs at the peak of cerebellar slow oscillation cycles. To probe this question, we calculated phase locking of the detected hippocampal SWR relative to the phase of cerebellar delta oscillations (<4 Hz, Figure 5D-F). We confirmed that hippocampal SWRs were phase locked to the delta rhythm observed in all three recorded cerebellar regions but that the strength of phase-locking was significantly greatest for Lob VI (Figure 5E, F). Given recent description of SWR propagation from hippocampus to neocortex (Nitzan et al., 2020), we also calculated SWR triggered spectral analysis of the cerebellar LFP (Supplemental Figure 4), which revealed no changes in the SWR frequency range (~150 Hz) and thus suggests hippocampal SWRs do not propagate directly or indirectly to the cerebellum. Finally, we investigated potential interplay between PSPs recorded in the cerebellum and hippocampal SWRs. To do so, we computed the probability of detecting a SWR following the detection of a PSP. We found a consistent decrease in SWR probability ~10 ms after the PSP peak in all cerebellar regions, which lasted for ~1 sec (compared with a shuffled dataset; Figure 5G). This reduction in SWR occurrence immediately following the PSPs suggests that the latter have an impact on local hippocampal activity.

## Discussion

Despite the extensive literature on sleep-related neurophysiological processes in the hippocampus (for review, see Klinzing et al., 2019) and recent advances in understanding of sleep-related processes in the cerebellum (Canto et al., 2017; Xu et al., 2020; Zhang et al., 2020), no report exists on potential functional or physiological interplay between these regions during sleep. Therefore, to bridge this knowledge gap, we investigated cerebello-hippocampal interactions as well as intrinsic cerebellar LFP dynamics across sleep states in mice.

### Intra-cerebellar LFP dynamics during sleep

We first profiled sleep-related activity within three distinct regions of the cerebellar cortex: Lob VI, Lob II/III and Crus I. We found modulation of cerebellar LFP activity across sleep states, both locally, in spatially segregated cerebellar lobules, and globally, in a coordinated manner across the cerebellar cortex. Overall, during REM and non-REM sleep, prominent delta (< 4 Hz) and VHFOs (~250 Hz) dominate the spectral profile in all three regions. The presence of VHFOs has been described previously in vermal lobules V/VIa of anaesthetised and awake, head-fixed rats, and attributed to activity in recurrent Purkinje cell collaterals (de Solages et al., 2008). In contrast, previous studies in freely moving mice have predominantly described high frequency oscillations in the cerebellar cortex (~150 Hz: Cheron et al., 2005, 2004; Servais et al., 2005), which is consistent with our observations during wake. Thus, given the high presence of VHFOs in head-fixed or sleeping animals they may be related to the absence of voluntary movements. Our findings additionally extend the work of de Solages et al. (2008) by showing for the first time that VHFOs are concurrently present across multiple cerebellar cortical regions, highly prevalent during sleep (above wake levels) and, furthermore, that they are temporally coordinated by local cerebellar delta rhythms during REM sleep.

During non-REM sleep, our description of prominent delta oscillations in the cerebellar cortex is in keeping with previous work showing that neocortical slow oscillations can entrain those in the cerebellar cortex (Roš et al., 2009a; Rowland et al., 2010; Xu et al., 2020). In contrast, REM sleep is traditionally associated with theta (~6-12 Hz) frequency, LFP oscillations. Our recordings from the cerebellum revealed no discernible increase in theta power during REM. Surprisingly, however, we did observe highly synchronous, large delta (< 4 Hz) oscillations in all three recorded cerebellar regions during REM. This may be a local circuity peculiarity; however, recent evidence rodent and human studies suggest that localized delta oscillations should be considered an integral component of REM sleep (Bernardi et al., 2019; Funk et al., 2016; Siclari and Tononi, 2017).

### Cerebello-hippocampal coherence during sleep

Next, we sought evidence for putative functional coupling, as measured by LFP-LFP coherence, between the hippocampus and cerebellum across sleep states. Of particular note, we found that hippocampus-Lob VI coherence was significantly and dynamically modulated across sleep states. Specifically, during non-REM, hippocampus-Lob VI delta coherence was significantly higher than during wake/REM whilst during REM, hippocampus-Lob VI theta coherence was significantly higher than during non-REM and, surprisingly, was also significantly modulated to above wake levels. The comparatively heightened, frequency-specific, offline coupling between hippocampus and Lob VI during non-REM illustrates the potential for both sleep stage and regionally specific interactions between the two brain structures in addition to those we previously described during active movement in the homecage (Watson et al., 2019). Indeed, this interaction may be supported by previously described anatomical connections linking Lob VI and hippocampus via the medial septum or the supramammillary nucleus (Watson et al., 2019).

### Putative PGO waves provide a potential substrate for cerebello-hippocampal interactions during REM

In addition to the ongoing LFP oscillations observed in the cerebello-hippocampal network during sleep, we also detected high amplitude PSPs in the cerebellar cortical recordings that were particularly abundant during REM sleep. Previous studies have described similar phasic events in the cerebellum and attributed them to propagation of PGO waves (Farber et al., 1980; Velluti et al., 1985). In keeping with our observations, PGO waves are found to be highly concentrated in REM epochs across multiple brain regions (Harlay et al., 1974; Marks et al., 1980; Pellet and Harley, 1977; Tsunematsu et al., 2020). Furthermore, PGO-waves are known to phase-lock to hippocampal theta rhythms during REM and modulate its frequency (increasing the preferred theta frequency; Karashima et al., 2007, 2002). We confirmed that this was also the case with phase-locking of cerebellar PSPs to the hippocampal LFP theta oscillation. Additionally, we also show that cerebellar PSPs are significantly phase-locked to the cerebellar delta oscillation cycle and cross-frequency analysis revealed that cerebellar delta modulation of hippocampal theta oscillations occurs preferentially when PSP content is high. Thus, overall, the most parsimonious explanation is that the PSPs we recorded in the cerebellum are propagated PGO waves. During REM, PGOs have been shown to play a crucial role in coordinating long range network dynamics underlying sleep-dependent cognitive processes required for establishment of fear memory (Datta, 2000; Datta et al., 1998; Datta and O’Malley, 2013). It is tempting to speculate that the PSPs observed in the cerebellum may play a similar role by coordinating interactions with the hippocampus and that this could subserve sleep-dependent memory formation of the explored environment.

### Sharp wave ripples link hippocampus and cerebellum during non-REM sleep

Sharp wave ripples (SWRs) are a prominent physiological feature of hippocampal activity during non-REM sleep. SWRs are fast oscillations during which both CA3 and CA1 pyramidal cells fire synchronously (Buzsáki, 1986; Csicsvari et al., 2000). Their occurrence is coordinated with cortical spindles, which are themselves synchronized with cortical slow-delta oscillations (Siapas and Wilson, 1998) and this tripartite interaction is important for memory formation (Girardeau et al., 2009; Latchoumane et al., 2017; Maingret et al., 2016). Consequently, we next examined the relationship between hippocampal SWRs and cerebellar LFP oscillations. SWR-triggered cerebellar LFP waveform averages in combination with phase-locking analysis revealed two main findings. Firstly, ERPs are present in the cerebellar cortical LFP at short latency (particularly in Lob VI) following hippocampal SWR onset. Previously, hippocampal SWRs have been shown to drive evoked LFP and single-unit changes in the cingulate cortex of a similar latency to those triggered by direct hippocampal electrical stimulation; as such SWR activity may be considered indicative of efferent flow from the hippocampal formation (Wang and Ikemoto, 2016). Indeed, we found that SWRs triggered cerebellar ERPs with a short onset latency of ~ 9.17 ms, which is in agreement with previous reports in anaesthetised rats (Saint-Cyr and Woodward, 1980; Saint-Cyr and Woodward, 1980b) in which direct electrical stimulation of the hippocampal fornix elicited both short latency mossy fibre (5-10 ms, routed via the pontine nuclei) and longer latency, climbing fibre (10-20 ms, routed via the inferior olive) responses within the cerebellar cortex. Also consistent with our results, topographical mapping of cerebellar responses following direct hippocampal electrical stimulation revealed evoked activity mainly in the Lob VI region of both cats (Newman and Reza, 1979) and rats (Saint-Cyr and Woodward, 1980b). Given the short latency of evoked SWR ERPs, particularly in Lob VI, it seems likely that SWR mediated hippocampal input to the cerebellum during non-REM sleep could be routed via the pontine nuclei. This hypothesis is further strengthened by evidence of a direct hippocampal projection to the pons (Schmahmann and Pandya, 1997).

Our second main finding was that SWRs are significantly phase locked to the up-state of cerebellar delta oscillations in striking similarity to the well described locking of SWRs to neocortical delta, which is thought to facilitate transfer of memory information from subcortical to cortical loci (Maingret et al., 2016). As cerebellar Purkinje cells are known to toggle between depolarizing ‘up-states’ and hyperpolarizing ‘down-states’ (see Engbers et al., 2013 for review) it may be considered that, during non-REM sleep, cerebellar delta oscillations may facilitate the transfer of hippocampal information via phase-locking of SWRs to periods of high Purkinje cell excitability thus favouring plasticity mechanisms associated with memory formation. Consistent with our finding that hippocampus - Lob VI delta coherence during non-REM sleep is higher compared to other cerebellar regions, SWRs were significantly more phase locked to Lob VI delta compared to Crus I or Lob II/III, further supporting the potential for regionally and sleep stage specific hippocampal-cerebellar interactions. Another physiological explanation of SWR links to cerebellar delta oscillations may be provided by the fact that SWRs are known to nest to the upstate of neocortical slow-delta oscillations. In turn, these neocortical oscillations have been shown to entrain cerebellar activity in both anaesthetised rats (Roš et al., 2009b; Rowland et al., 2010; Schwarz, 2010) and naturally sleeping monkeys (Xu et al., 2020). Thus, SWR phase-locking to cerebellar delta oscillations may be mediated via upstream SWR-to-neocortical delta nesting.

Taken together, the SWR-triggered evoked LFP activity recorded in the cerebellar cortex and their coordination with cerebellar delta oscillations illustrate the existence of robust physiological links and candidate mechanisms that could subserve hippocampal-cerebellar (particularly Lob VI) interactions during non-REM sleep.

In summary, our findings illustrate the presence of multiple physiological events in the cerebello-hippocampal network during sleep (Supplemental Figure 5), centred around prominent delta oscillations. In particular, we have highlighted lobule specific hippocampus-to-cerebellum directed interaction during non-REM sleep mediated via SWR and delta oscillations. During REM, we identified prominent cerebellar delta oscillations and associated PSPs, which modulate both local cerebellar VHFO and distant hippocampal theta oscillations. Thus, it appears that delta oscillations play a key role in temporal coordination both within and between these regions. Similar sleep-stage specific, transient physiological events have been shown to be important for memory formation across other brain circuits, and by analogy it may be hypothesised that the events described in the hippocampal-cerebellar network serve a similar role in affording the two regions the ability to preferentially interact during windows of enhanced synaptic plasticity.

As the cerebellum is thought to compute forward models that predict sensory consequences of actions adapted to a particular context, we speculate that the aforementioned mechanisms may allow offline updating and refinement of such models across the cerebello-hippocampal network during sleep. These refinements may be important during online navigation behaviours in which animals must predict the spatial consequences of their movements.

## Material and Methods

### Mice

20 adult male C57BL6-J mice were used for this study (Janvier, France). Mice were housed individually under a 12 hr light/12 hr dark cycle (light cycle beginning at 8 am) and received food and water *ad libitum*. All behavioural experiments were performed in accordance with the official European guidelines for the care and use of laboratory animals (86/609/EEC) and in accordance with the Policies of the French Committee of Ethics (Decrees n° 87-848 and n° 2001-424). The animal housing facility of the laboratory where experiments were made is fully accredited by the French Direction of Veterinary Services (B-75-05-24, 18 May 2010). The protocol was approved by the Committee on the Ethics of Animal Experiments (APAFIS#4315-2016042708195884v1). Data obtained from the mice used in this study have been published in Watson et al., (2019).

### Surgery

Surgical and implant procedures have been already described in Watson et al., (2019). Briefly, surgical implantation was performed under constant isoflurane anaesthesia (1.5 %) combined with oxygen (1.5 L/min). Animals were placed in a stereotaxic frame device (David Kopf Instruments, USA) and an incision was performed in order to expose the scalp. Coordinates for implantation were calculated from bregma following the references given in Franklin and Paxinos, 2007. We targeted bilateral hippocampi (AP −2.2 mm, ML ± 2.0 mm, DV 1.0 mm), cerebellar lobules II/III (AP −5.52 mm, ML 0 mm, DV 1.8 mm), cerebellar lobule VI (AP −6.72 mm, ML 0 mm, DV 0.1 mm) and left cerebellar Crus I (AP - 6.24 mm, ML 2.5 mm, DV 0.1 mm). Small craniotomies were performed over the target regions using a drill and the dura was carefully removed with a needle. Two wires of 140 μm diameter teflon coated stainless-steel (A-M system, USA) were twisted together to create bipolar LFP recording electrodes (interpolar distance ~0.5 mm) and were implanted in the brain. Pairs of flexible stainless-steel wires were also sutured to the neck muscles to obtain EMG recordings (Cooner wire, USA). 14 mice were also implanted with bipolar stimulation electrodes (same as the LFP electrodes but with interpolar distance of ~140 μm) in the left medial forebrain bundle (AP −1.4 mm, ML 1.2 mm, DV 4.8 mm) to serve as a reward signal in a set of experiments already published (Watson et al., 2019). All electrodes were attached to an electrode interface board (EIB-18, Neuralynx, USA) and the assembly was fixed to the skull using a combination of UV activated cement (SpeedCem, Henry Shein, UK), SuperBond (SunMedical, Japan) and dental cement (Simplex Rapid, Kemdent, UK). Four miniature screws (Antrin, USA) were also attached to the skull for additional support and to serve as recording ground. Animals were given a minimum of 5 days post-surgery recovery time before experiments commenced.

### Electrophysiological recordings

The EMG and LFP recordings were obtained via a unity-gain headstage preamplifier (HS-18, Neuralynx, USA) and a Digital Lynx SX system and Cheetah software (Neuralynx, USA). Signals were bandpass-filtered between 0.1 and 600 Hz and sampled at 1 kHz. Mouse position was tracked at 30 Hz using video tracker software and infra-red LEDs attached to the headstage (Neuralynx, USA).

### Histology

After completion of the experiments, mice were deeply anaesthetized with ketamine/xylazine solution (150 mg/Kg) and electrolytic lesions were created by passing a positive current through the electrodes (30 μA, 10 s). The animals were then perfused transcardially with saline (0.9 %) followed by paraformaldehyde (4%). Once perfused, the electrode assembly was carefully removed, the brain extracted and post-fixed in paraformaldehyde (4%) for 24 hr and then embedded in agarose (3%). 50 μm sagittal and coronal sections for the cerebellum and the hippocampus, respectively, were made using a vibratome. The sections were mounted on gelatinized slides and stained with cresyl violet. The electrolytic lesions were then identified in order to reconstruct the recording locations using standard maps with reference to a stereotaxic atlas (Franklin and Paxinos, 2007). The anatomical location of the electrodes used in this study were verified in Watson et al., (2019).

### Behaviour and sleep scoring

The animals were recorded during the day (between 10 am and 6 pm) in their home-cages (30 cm x 10 cm x 10 cm), with the lid removed and the lights off for periods up to 4 hr. Recordings were made prior to training in a linear track task (described in Watson et al., 2019). The homecage environment was familiar to the mice as they had been housed in it since completion of implantation surgery. Animals exhibited different behavioural states that were scored off-line and included in four categories: active wakefulness, resting, non-REM sleep and REM sleep. The scoring was semi-automatically performed using multi-parameter thresholds based upon instantaneous speed, neck EMG (rectified, smoothed with 1 s window and z-scored) and theta/delta ratio in the hippocampal LFP (z-scored and calculated in 100 ms bins). Thus, epochs of more than 4 s with instantaneous speed above 3 cm/s were considered as active wakefulness, periods of less than 30 s where the speed stayed below 3 cm/s and were surrounded by active wakefulness epochs were considered as rest. Similarly, when the EMG recording was optimal, a manually selected threshold was also used to discriminate between rest and sleep epochs. We finally used manual thresholding of the theta/delta ratio to separate non-REM and REM sleep epochs. Only epochs of more than four seconds were selected for further analysis.

### Data preprocessing

All the data processing was performed using custom-made MATLAB scripts (Mathworks, USA). Raw signals were pre-processed by applying a notch filter to remove electrical line noise (filter centred to 50 Hz). Voltages were z-scored to reduce overall differences in amplitude in the signal recorded from different electrodes.

### Offline detection of ripples, slow oscillations and phasic events

Discrete sharp-waves ripples (SWRs) were detected using criteria employed elsewhere (Maingret et al., 2016). The raw signal was then filtered using a second order zero-phase bandpass filter between 150-250 Hz, squared, smoothed (using 8 ms running average) and z-scored. The SWRs were defined as events in which the transformed signal remained above 2 z-scores for 30 to 100 ms with a peak above 5 z-scores.

Slow oscillations in the cerebellum were divided into individual cycles which were identified as follows: first the raw signal was filtered using a second order zero-phase bandpass filter between 0-1-4Hz. The instantaneous phase was obtained by applying a Hilbert transform to the filtered signal. Each cycle was defined by the epoch between two negative peaks on the cosine of the instantaneous phase. Only cycles which lasted more than 1 s and presented peak amplitudes higher than 4 z-scores and trough amplitudes no lower than 2 z-scores were further analysed.

For the detection of PSPs (putative p-waves) in the cerebellum, we first filtered the signal using a second order zero-phase bandpass filter between 5-80 Hz, squared it, and then calculated the z-score across all obtained sleep epochs. A double threshold strategy was then applied, first to the transformed signal so that epochs separated by at least 75 ms with values above 1 z-score and a peak in the filtered data above 4 z-scores were considered as events.

### Spectral analysis

All the spectral analyses were performed employing freely available signal processing toolboxes. For computing the overall spectral power and spectral coherence across sleep-states, a multi-taper Fourier transform (Chronux toolbox) was computed with 4 s sliding windows in 1 s steps and using 4 tapers. The mean power spectrum and coherence was obtained for each behavioural state by averaging the spectrogram and coherogram from the identified epochs. To reduce the impact of the different data durations for each state, we split the epochs to match with the minimal data duration and then we averaged them. For computing the triggered spectrogram and coherograms we used the continuous wavelet transform and wavelet coherence (wavelet toolbox, MATLAB). We used analytic Morse wavelets with 8 octaves and 48 voices per octave.

### Phase locking of phasic events and slow-wave triggered power spectra and coherograms

Cerebellar and hippocampal signals were filtered using a second order zero-phase bandpass filter between 0.1 - 4 Hz and 6 - 12 Hz, respectively. The instantaneous phase of the cerebellar infra-slow oscillations and the hippocampal theta oscillations were obtained by applying a Hilbert transform to the filtered signals. The phase locking of the detected cerebellar phasic events to these rhythms was tested using circular statistics (Matlab) and significance was determined as Raleigh test p-value < 0.05.

To compute the cerebellar slow-wave triggered spectrogram and coherogram, for each cerebellar slow-wave, the obtained time-frequency series was matched with the corresponding instantaneous phase of the cerebellar delta oscillations (0.1-4 Hz) and then averaged by phase bins of 7.35 °.

### Cross-correlation between PSPs and SWRs

The timing of the SWRs was calculated relative to the time of the peak of the detected PSPs. After binning at 25 ms resolution, the probability density function (count normalized by the total number of elements multiplied by the bin width) was obtained and smoothed (using a 250 ms running average). Statistical significance of the obtained cross-correlation was performed by artificially generating PSP trains of similar density as observed in the actual data, drawn at random from a Poisson distribution, and recomputing the probability density function.

### Statistical analysis

Statistical analyses were conducted using MATLAB Statistical Toolbox and Prism (Graphpad, USA). Normality was assessed using a Shapiro-Wilk test. Parametric and non-parametric tests were then used accordingly. Paired analyses were employed when possible.

## Author Contributions

Arturo Torres-Herraez, Software, Formal analysis, Validation, Investigation, Visualization, Methodology, Writing-original draft; Thomas Charles Watson, Conceptualization, Supervision, Formal analysis, Validation, Visualization, Investigation, Methodology, Writing-original draft, review and editing; Laure Rondi-Reig, Conceptualization, Resources, Supervision, Funding acquisition, Methodology, Writing-critical revision, review and editing, Project administration.

## Acknowledgments

This work was supported by the Fondation pour la Recherche Médicale DEQ20160334907-France, the National Agency for Research ANR-17-CE16-0019-SynPredict, CNRS, Inserm and Sorbonne University (LRR). We thank all members of the CEZAME team for helpful discussions of the experiments and manuscript. We gratefully acknowledge the IBPS animal facility staff for their support.

## Supplementary Figures

**Supplemental Figure 1.**
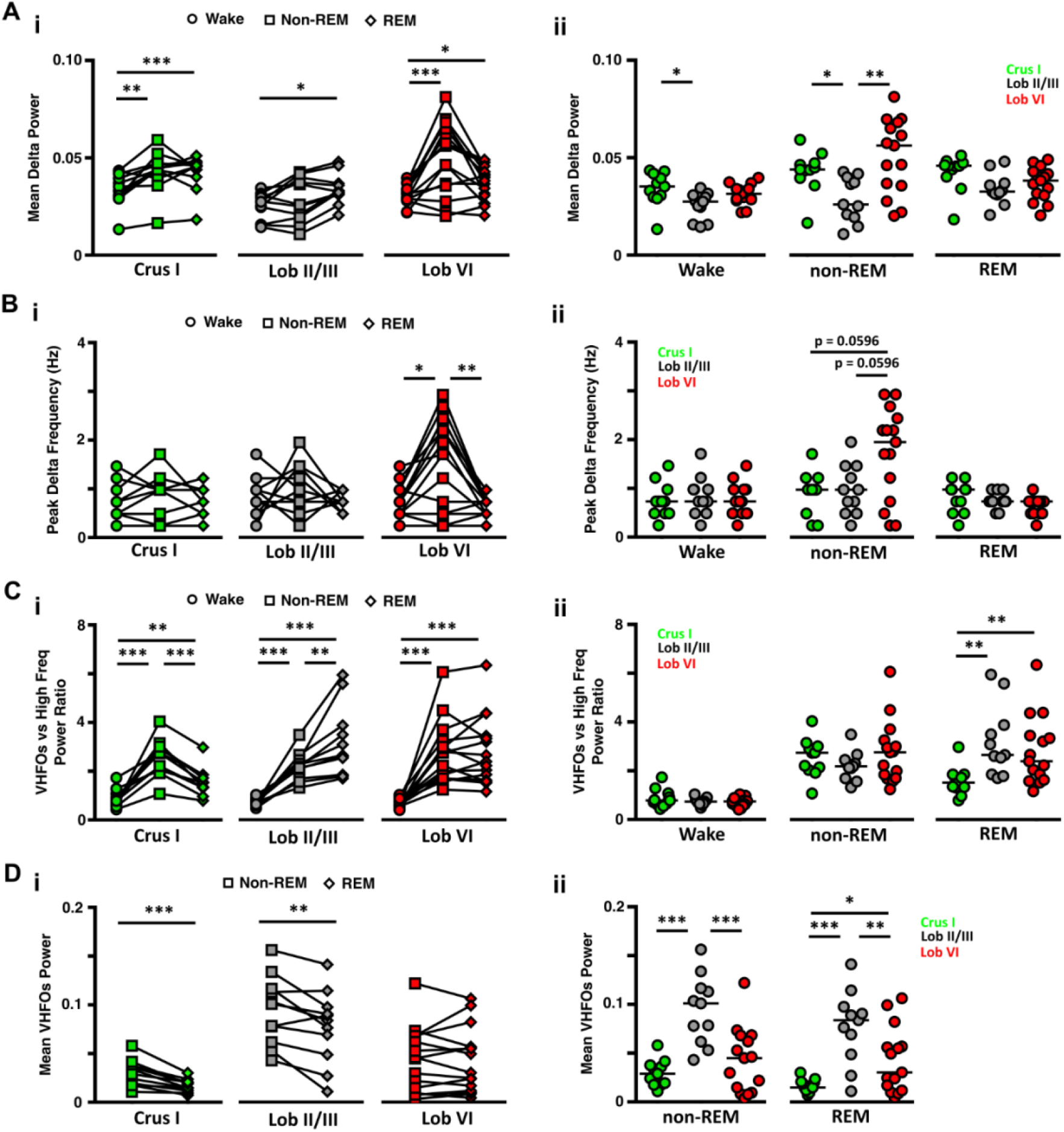
LFP power in delta (0.1-4 Hz Hz) and VHFO (240-280 Hz) frequency bands during wake and sleep. **A, i,** The power of delta oscillations varied between wake and the different sleep states within the three cerebellar regions (Crus I, n = 11, Friedman test, p = 0.0004, multiple comparisons with FDR correction: awake vs non-REM, p = 0.0015, awake vs REM, p = 0.0007, non-REM vs REM, p = 0.2344; Lob II/III, n = 11, Friedman test, p = 0.0273, multiple comparisons with FDR correction: awake vs non-REM, p = 0.3657, awake vs REM, p = 0.0221, non-REM vs REM, p = 0.0578; Lob VI, n = 15, Friedman test, p = 0.0023, multiple comparisons with FDR correction: awake vs non-REM, p = 0.0011, awake vs REM, p = 0.0468, non-REM vs REM, p = 0.1009). **ii**, Similarly, differences between cerebellar regions were observed both during wake (Kruskal-Wallis test, p = 0.0133, multiple comparisons with FDR correction: Crus I vs Lob II/III, p = 0.0075, Crus I vs Lob VI, p = 0.1473, Lob II/III vs Lob VI, p = 0.0633) and non-REM sleep (Kruskal-Wallis test, p = 0.0040, multiple comparisons with FDR correction: Crus I vs Lob II/III, p = 0.0117, Crus I vs Lob VI, p = 0.1466, Lob II/III vs Lob VI, p = 0.0012), but not during REM (Kruskal-Wallis test, p = 0.0540). **B**, **i**, The peak delta band frequency differed across sleep states only in Lob VI (Crus I, Friedman test, p = 0.1322; Lob II/III, Friedman test, p = 0.2233; Lob VI, Friedman test, p = 0.009, multiple comparisons with FDR correction: awake vs non-REM, p = 0.0107, awake vs REM, p = 0.1955, non-REM vs REM, p = 0.0039). **ii**, Differences in the peak delta frequency band across cerebellar regions were also restricted to non-REM epochs (wake, Kruskal Wallis test, p = 0.7732, non-REM, Kruskal Wallis test, p = 0.0364, multiple comparisons with FDR correction: Crus I vs Lob II/III, p = 0.9095, Crus I vs Lob VI, p = 0.0596, Lob II/III vs Lob VI, p = 0.0596). **C**, **i**, During wake, the power of VHFOs was lower compared to high frequency oscillations in all regions, as indicated by the mean values below 1. In contrast, during sleep, across all recorded cerebellar regions, the ratio of VFHO to high frequency oscillation power increased significantly compared to wake (shifted to values >1), and also differed between sleep states (Crus I, repeated measures ANOVA, p < 0.0001, multiple comparisons with FDR correction: awake vs non-REM, p < 0.0001, awake vs REM, p = 0.0046, non-REM vs REM, p < 0.0001; Lob II/III, repeated measures ANOVA, p = 0.0002, multiple comparisons with FDR correction: wake vs non-REM, p < 0.0001, wake vs REM, p = 0.0003, non-REM vs REM, p = 0.0071; Lob VI, repeated measures ANOVA, p < 0.0001, multiple comparisons with FDR correction: wake vs non-REM, p < 0.0001, wake vs REM, p < 0.0001, non-REM vs REM, p = 0.7226). **ii**, Significant differences across cerebellar regions were restricted to REM epochs when Crus I shown significantly smaller ratios than the other two regions (wake, one-way ANOVA, p = 0.2252, non-REM, one-way ANOVA, p = 0.3705, REM, one-way ANOVA, p = 0.0098, multiple comparisons with FDR correction, Crus I vs Lob II/III, p = 0.0042, Crus I vs Lob VI, p = 0.0083, Lob II/III vs Lob VI, p = 0.1538). **D**, **i**, VHFO power was significantly reduced during REM sleep compared with non-REM in Crus I (Paired t-test, p = 0.0005) and Lob II/III (Paired t-test, p = 0.0067) but remained unchanged in Lob VI (Paired t-test, p = 0.9137). **ii**, VHFOs power was significantly higher in Lob II during both non-REM (one-way ANOVA, p < 0.0001, multiple comparisons with FDR correction, Crus I vs Lob II/III, p < 0.0001, Crus I vs Lob VI, p = 0.1020, Lob II/III vs Lob VI, p < 0.0001) and REM sleep (one-way ANOVA, p = 0.0002, multiple comparisons with FDR correction, Crus I vs Lob II/III, p = 0.0001, Crus I vs Lob VI, p = 0.0393, Lob II/III vs Lob VI, p = 0.0092).

**Supplemental Figure 2.**
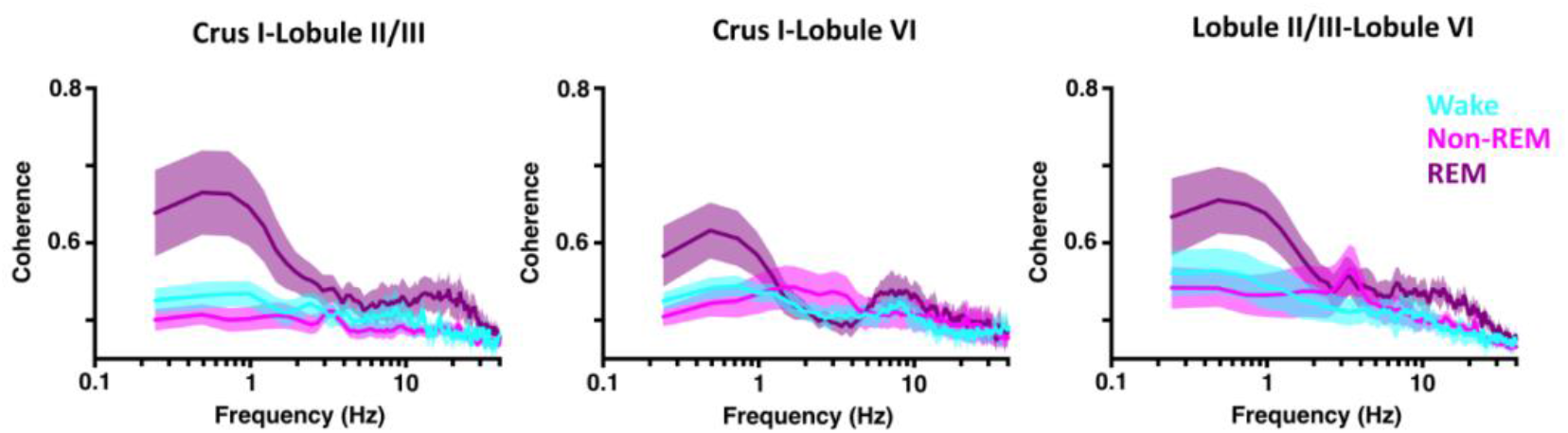
Intra-cerebellar delta coherence is highest during REM sleep. Delta band (< 4Hz) coherence was elevated during REM sleep compared with both awake and non-REM in all combinations of cerebellar recordings (Crus I-Lob II/III, n = 8; Crus I-Lob VI, n = 7; Lob II/III-Lob VI, n = 7).

**Supplemental Figure 3.**
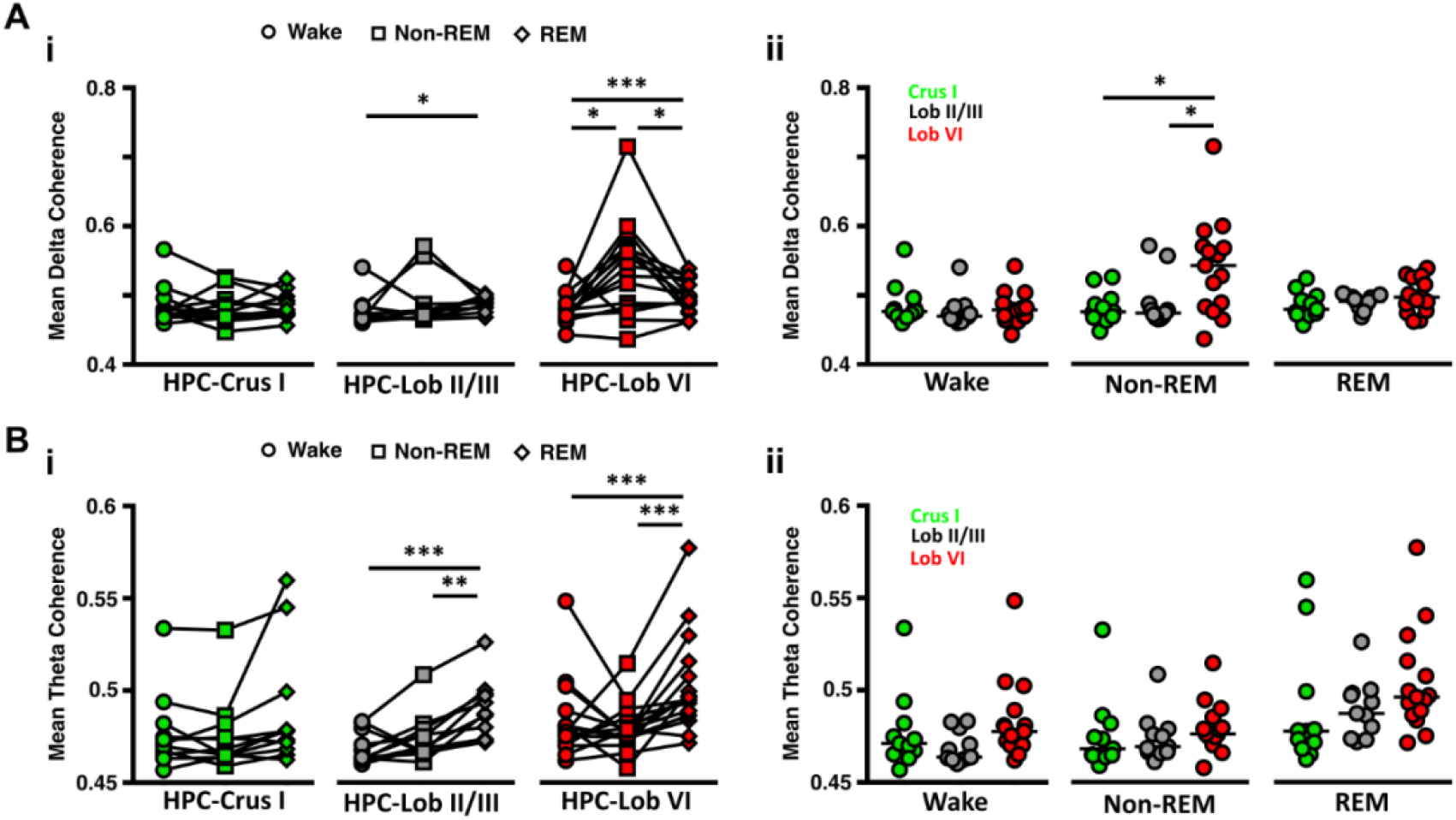
Cerebello-hippocampal coherence in the delta (<4 Hz) and theta (6-12 Hz) frequency ranges is modulated across sleep-states. **A**, **i**, Significant modulation across sleep states in delta coherence was found between hippocampus and Lob II/III (n = 11, Friedman test, p = 0.0273, multiple comparisons with FDR correction: wake vs non-REM p = 0.3657, wake vs REM p = 0.0221, non-REM vs REM p = 0.0578), and particularly between the hippocampus and Lob VI (n = 15, Friedman test, p = 0.0006, multiple comparisons with FDR correction: wake vs non-REM p = 0.0001, wake vs REM p = 0.0149, non-REM vs REM p = 0.0351). Hippocampus-Crus I delta coherence remained unchanged (n = 11, Friedman test, p = 0.8438). **ii**, Significant differences in delta coherence between the cerebello-hippocampal combinations were restricted to non-REM epochs, with HPC-Lob VI levels being significantly higher than other combinations (wake, Kruskal-Wallis test, p = 0.4283; non-REM, Kruskal-Wallis test, p = 0.0225, multiple comparisons with FDR correction: HPC-Crus I vs HPC-Lob II/III, p = 0.3225, HPC-Crus I vs HPC-Lob VI, p = 0.0122, HPC-Lob II/III vs HPC-Lob VI, p = 0.0122; REM, Kruskal-Wallis test, p = 0.2826). **B**, **i**, Similarly, theta coherence was significantly increased during REM sleep between HPC-Lob II/III (n = 11, Friedman test, p = 0.0002, multiple comparisons with FDR correction: wake vs non-REM p = 0.1378, wake vs REM p = 0.0003, non-REM vs REM p = 0.0029) and HPC-Lob VI (n = 15, Friedman test, p = 0.0007, multiple comparisons with FDR correction: wake vs non-REM p = 0.2503, wake vs REM p = 0.001, non-REM vs REM p = 0.0005), but remained stable across sleep states between the HPC-Crus I (n = 11, Friedman test, p = 0.1632). **ii,** In contrast, no differences across cerebello-hipocampal combinations were found during any of the states analysed (wake, Kruskal-Wallis test, p = 0.0677; non-REM, Kruskal-Wallis test, p = 0.1994; REM, Kruskal-Wallis test, p = 0.0979).

**Supplemental Figure 4.**
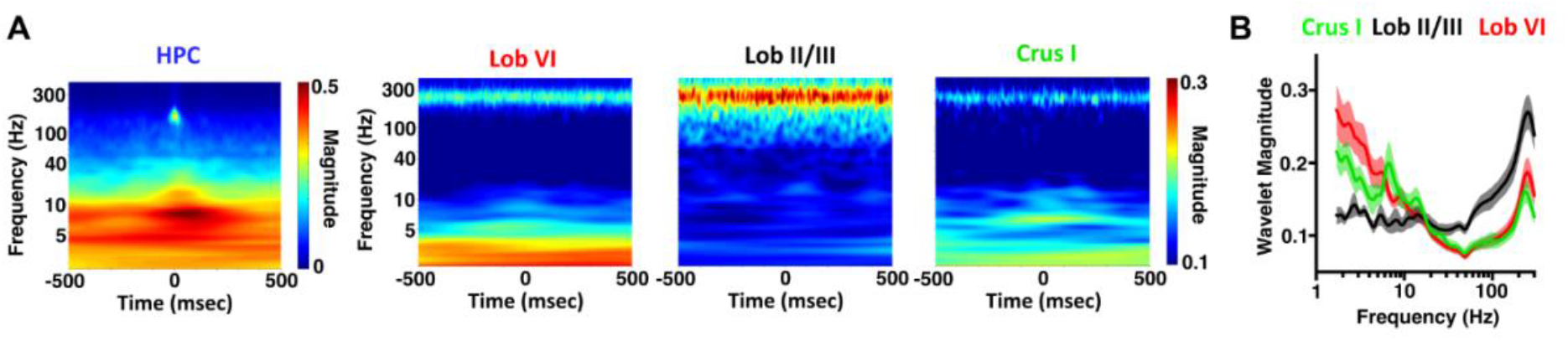
SWR triggered spectral analysis. **A**, Averaged power spectra triggered by detected hippocampal SWR events (time zero). Nitzan et al., (2020) have recently shown that SWRs can propagate to the cortex. Here we observe discrete SWR activity in the hippocampal spectrogram (~150Hz) that is not present in the cerebellum. **B**, Mean cerebellar power spectra calculated during the 500ms following ripple detection.

**Supplemental Figure 5.**
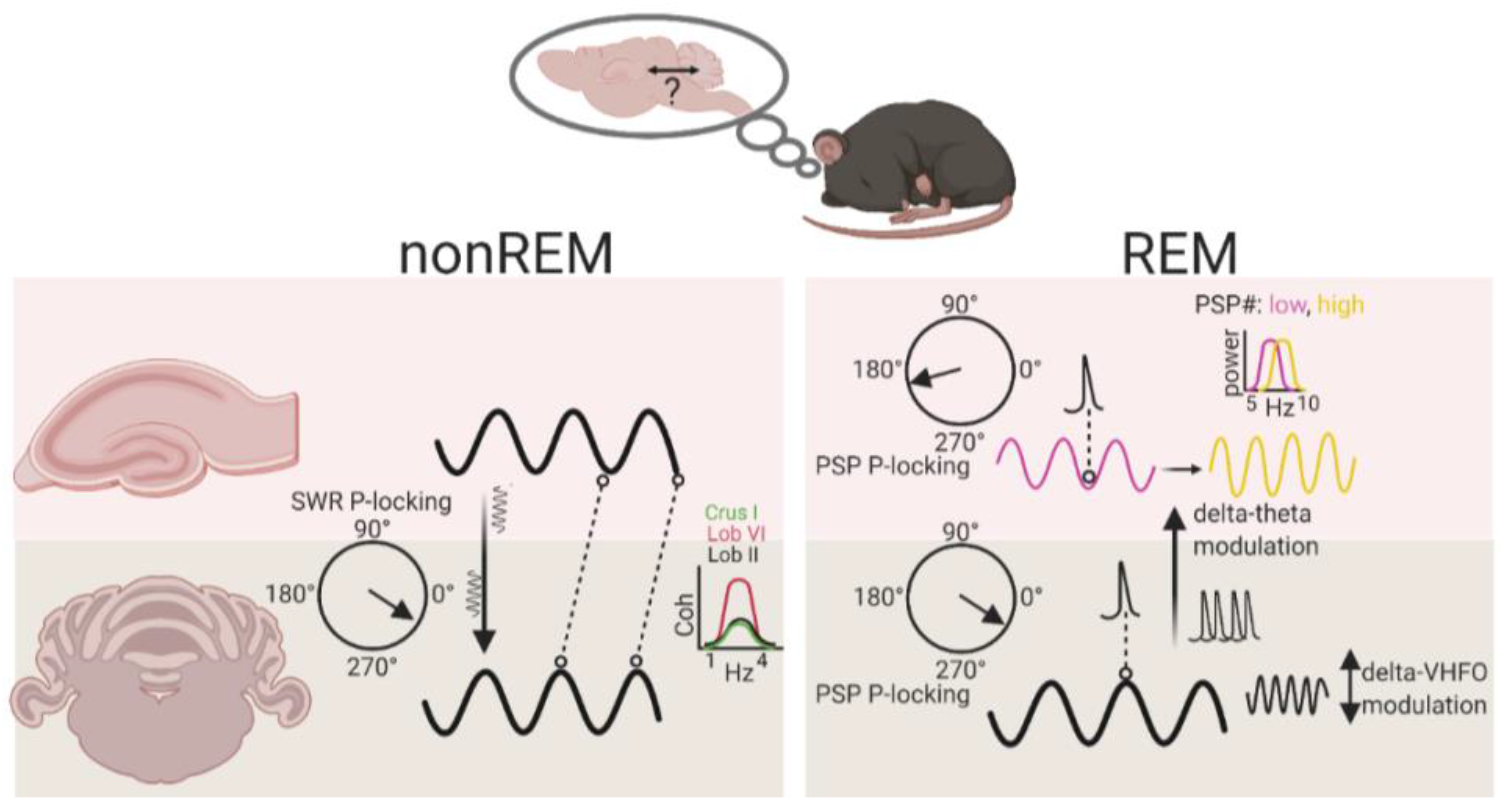
Summary diagram of main findings illustrating sleep-stage specific physiological events and interactions within the cerebello-hippocampal network. During nonREM, coherence within the delta frequency range is highest between hippocampus and lobule VI compared to other cerebellar lobules. In addition, hippocampal sharp wave ripples (SWR) are phase locked to the cerebellar delta oscillation (black line) and drive modulation of cerebellar LFP activity. During REM, PSPs are significantly phase locked to both the trough of hippocampal theta (purple line) and the peak of cerebellar delta oscillations (black line). Within the cerebellum, delta oscillations can modulate activity within the VHFO range during REM. Additionally, cerebellar delta oscillations and associated PSPs modulate the frequency of hippocampal theta rhythms (from ~ 7.5 Hz (low, purple line) to 8 Hz (high, yellow line)). Abbreviations: SWR, sharp wave ripple; P-locking, phase locking; PSP, phasic sharp potential; VHFO, very high frequency oscillation. Created with BioRender.com.

